# Maize phytocytokines and microbial-patterns trigger antagonistic features in co-incidence with wounding and fungal pathogens

**DOI:** 10.1101/2022.12.21.521402

**Authors:** Maurice Koenig, Daniel Moser, Julian Leusner, Jasper Depotter, Gunther Doehlemann, Johana Misas Villamil

## Abstract

Phytocytokines are signalling peptides that activate immune responses and alarm bystander cells of danger. The pathways of phytocytokine perception and activation are thought to be shared between exogenous danger signals such as microbe-associated molecular patterns (MAMPs) and endogenous, passively released, damage-associated molecular patterns (DAMPs). However, downstream responses triggered by danger molecules and their effect in plant survival is still largely unknown. Here, we have identified three biologically active maize orthologues of phytocytokines previously described in other plants. The maize phytocytokines show common features with MAMPs, including the induction of immune related genes and activation of papain-like cysteine proteases. In contrast to MAMPs, the phytocytokines do not promote cell death in the presence of wounding. In infection assays with fungal pathogens of two different life styles we found that phytocytokines affect the development of disease symptoms, likely due to the activation of phytohormonal pathways. Collectively, our results show that phytocytokines and MAMPs trigger unique and antagonistic features of immunity. We propose a model in which phytocytokines activate immune responses partially similar to MAMPs but in contrast to microbial signals, they act as danger and survival molecules to the surrounding cells. Future studies will focus on the components determining the divergence of signalling outputs upon phytocytokine activation.

## Introduction

To recognize microorganisms, plants have evolved a diverse range of pattern recognition receptors (PRR) that detect conserved patterns characteristic of an infection. These PRRs can detect microbe-associated molecular patterns (MAMPs) or damage-associated molecular patterns (DAMPs) (Peng et al., 2018; Saijo et al., 2018). MAMPs can be microbial proteins, polysaccharides, peptidoglycan, lipids, or other conserved molecules (Boller & Felix, 2009; Boller & Flury, 2012). A well-known MAMP is the conserved 22 amino acid peptide of bacterial flagellin (flg22), which is recognized by the leucine-rich repeat receptor-like kinase (LRR-RLK) FLAGELLIN SENSITIVE (FLS2) (Chinchilla et al., 2006). The detection of MAMPs triggers defenses responses leading to MAMP-triggered immunity (MTI) (Peng et al., 2018). MTI defense response includes an increase of cytosolic Ca^2+^, production of apoplastic reactive oxygen species (ROS) and mitogen-activated protein kinases (MAPK) cascade activation (Saijo et al., 2018). These signals are quickly followed by the synthesis of phytohormones such as ethylene (ETH) and salicylic acid (SA) (Boller & Felix, 2009). Finally, hormonal activation induces stomatal closure, callose deposition, as well as transcriptional and metabolic reprogramming (Boller & Felix, 2009; Saijo et al., 2018).

In contrast to MAMPs, DAMPs are plant derived peptides, sugars, extracellular DNA, or other molecules which are passively released upon leakage of damaged cells into the apoplast, or after degradation by microbial enzymes (Gust et al., 2017; Hou, Liu, et al., 2019; Li et al., 2020). Defense responses to DAMPs are thought to protect against mechanical or cellular damage triggering defense responses similar to MTI (Hou, Liu, et al., 2019). One example of a cell wall-derived DAMP are oligogalacturonides (OGs). These are fragments of the pectin polysaccharide homogalacturonan, a linear polymer of α-1-4–linked galacturonic acid, that assist in maintaining cell wall integrity (Pontiggia et al., 2020). OGs can be released mechanically or by pathogen-secreted hydrolytic enzymes and recognized by the receptor WAK1 (Brutus et al., 2010; Decreux & Messiaen, 2005; Denoux et al., 2008; Pontiggia et al., 2020).

Endogenous signaling peptides were previously classified as DAMPs but recently “danger signals” have been re-classified into molecules passively released (classical DAMPs) or as plant elicitor peptides processed or secreted upon “danger”, or phytocytokines (Gust et al., 2017; Luo, 2012). Phytocytokines are divided into two classes, depending on whether the precursor protein (propeptide) contains a signal peptide or not. The majority of phytocytokines identified to date belong to the secreted elicitor peptides (Hou et al., 2021; Li et al., 2020), including hydroxyproline-rich systemins (HypSys) (Chen et al., 2008), PAMP-induced secreted peptide 1 (PIP1)/PIP2 (Hou et al., 2014), serine-rich endogenous peptide 12 (SCOOP12) (Gully et al., 2019), phytosulfokines (PSKs) (Amano et al., 2007), plant peptide containing sulphated tyrosine 1 (PSY1) (Amano et al., 2007), inflorescence deficient in abscission (IDA)/IDA-LIKE 6 (IDL6) (Butenko et al., 2003), root meristem growth factors (RGFs)/GOLVENs (GLVs) (Stegmann et al., 2022), immune related peptide (IRP) (Wang et al., 2020), CAP-derived peptide (CAPE) (Chien et al., 2015) and rapid alkalization factors (RALFs) (Stegmann et al., 2017). On the contrary, Systemin (McGurl et al., 1992), plant elicitor peptides (PEPs) (Huffaker et al., 2006, 2011; Nakaminami et al., 2018; Yamaguchi et al., 2011) and the *Z. mays* immune signaling peptide 1 (Zip1) (Ziemann et al., 2018) belong to the non-secreted phytocytokines (Hou et al., 2021). In contrast to DAMPs, phytocytokines are quickly transcriptionally activated, produced as propeptides and then the signaling peptide is actively released by plant proteases during pathogen attack or wounding. For instance, the plant elicitor peptide 1 (PEP1) is released within minutes from its precursor, ProPEP1, upon wounding of plant cells (Huffaker et al., 2006, 2011). Wounding triggers Ca^2+^ accumulation in the cytosol activating the plant protease METACASPASE 4 (MC4) which releases PEP1 from the vacuolar membrane into the extracellular space, via an unknown mechanism, promoting the recognition by the bystander cells via the LRR-RLK cell surface receptors PEPR1 and PEPR2 (Hander et al., 2019; Yamaguchi et al., 2006). Besides their role as signaling molecules in immunity, phytocytokines can also modulate developmental processes and act as cell-to-cell communication signals (Rzemieniewski & Stegmann, 2022).

Conservation of phytocytokines across different plant species can be very low on sequence level and it is rarely studied on the functional level. Some phytocytokines such as RALFs and PEPs are ubiquitously found in many plant species (Gust et al., 2017). Some other phytocytokines have been identified in most species of a family such as Systemin only present in Solanaceae or IRP specific for Poacea (Li et al., 2020; Ryan & Pearce, 2003), but even some have been specifically found in one species such as Zip1 which is specific for *Zea mays* and its wild ancestor teosinte (Depotter et al., 2022). Evolutionary arms-race might have led to a sequence diversification of some phytocytokines and it remains largely unknown, if signaling peptides found in dicots also function similarly in monocots.

The plant response against exogenous and endogenous danger signals and the downstream responses, such as cell death provoked by recognition of these molecules is a central question of our study. Using a bioinformatics pipeline, we aimed to identify maize orthologues of phytocytokines described in other species to be involved in plant immunity. We then addressed the bioactivity of candidate peptides and identified three new maize phytocytokines with functions in immune signaling and pathogen resistance. We found that maize phytocytokines trigger overlapping signaling pathways with MAMPs, but they also show unique features, such as the arrest of cell death in the co-incidence of wounding. In summary, our study reveals that perception of phytocytokines and MAMPs lead to distinct outcomes in plant immunity. In contrast to MAMPS, which promote cell death in the presense of wounding, phytocytokines act as danger signals but do not contribute to regulated cell death.

## Results

### Mining the maize genome for small signaling peptides

A literature search was performed to identify signaling peptides that are involved in plant immunity. 18 immune-related propeptides were found in monocots and dicots, from which 10 from 18 sequences belong to *A. thaliana* (Table 1). The GOLVEN2 (GLV2) peptide (Stegmann et al., 2022) was published during preparation of this manuscript and thus it could not be included in our analysis. To identify putative peptide orthologues in maize, a database containing 78 predicted plant proteomes including 31 monocots, 43 dicots and 4 from other phyla was generated (Suppl. Table 1). A search in this database was performed using a position-specific iterative Basic Local Alignment Search Tool (psiBLAST) due to its sensitivity and capacity to find orthologues with a lower local sequence similarity (Altschul et al., 1997). These 78 proteomes were selected to generate a position-specific score matrix covering as much proteome diversity as possible for the psiBLAST search to identify orthologues between very distantly related species (e.g. between *A. thaliana* and *Z. mays*). In addition, a BLASTp search (Altschul et al., 1990) was performed to compare to the psiBLAST search results. After removal of low quality (protein identities < 10% and query coverage < 25%) and redundant entries, the best hit propeptide per organism was selected for further analysis, yielding in total 964 pre-filtered candidates (Suppl. Fig. 1). Notably, the psiBLAST searches yielded more hits compared to BLASTp searches (Suppl. Fig. 2), confirming that an iterative search can improve the sensitivity of a BLASTp to find distant orthologues. The majority of the psiBLAST searches yielded between 33 to 79 hits per queried propeptide (e. g. OsProIRP: 33, AtPrePIP1: 68, and AtProPEP1: 60, Suppl. Fig. 1, initial filtering). Interestingly, only few or no hits were found in searches for the propeptides of GmProPEP890, GmProPEP914, AtProSCOOP12 and ProZip1 (Suppl. Fig. 1, length filtering), indicating that orthologues of these propeptides are encoded only in a small number of genomes. To reduce the number of false-positive hits, sequences with low sequence length similarity to the query (1.5-time longer or shorter) were removed. This filtering affected mostly hits with sequence identities below 30%, especially hits for NtPreproHypSys and SlProsystemin (Suppl. Fig. 1, length filtering). Afterwards, all hits were aligned with CLUSTAL Omega (Goujon et al., 2010) and putative peptides were identified by alignment. The similarity to the queried peptide (motif score) was calculated based on the alignment of the discovered peptide to the queried peptide using a BLOSUM62 matrix (Henikoff & Henikoff, 1992) and the self-alignment of the queried peptides as reference to account for differing peptide lengths. Hits with a motif score below 10% were removed, since this was the scoring were ZmPEP1 was found via our pipeline with AtPEP1 as a query sequence (Suppl. Fig. 1, iterative motif filtering).

**Table 1:**
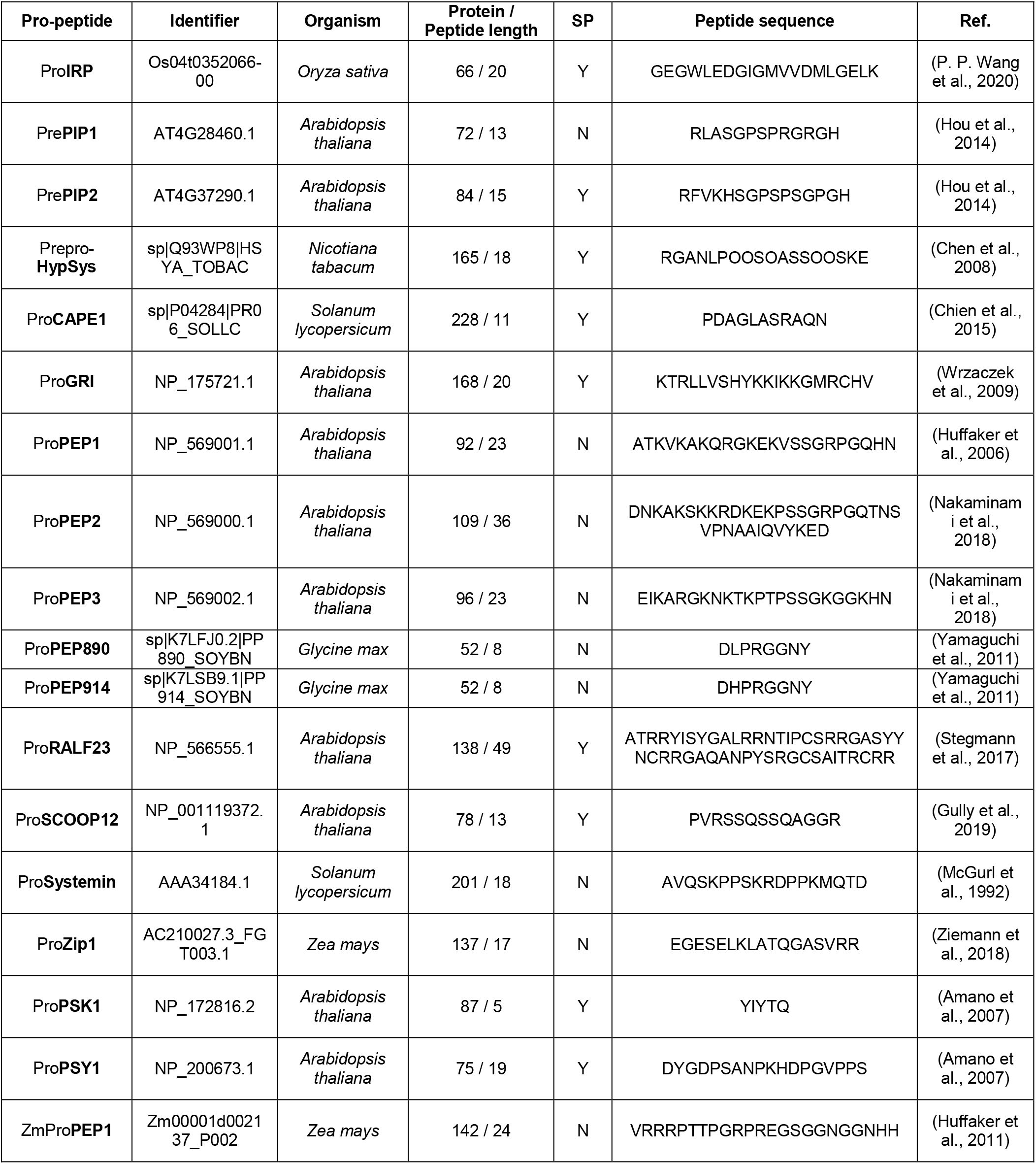
Propeptides and their properties used in the BLAST search. Identifier, source organism, protein / peptide amino acid sequence length, presence (Y) / absence (N) of signal peptide (SP), peptide sequence and literature reference (Ref.) where the query peptide was found are listed.

To compare whether certain peptide orthologues were restricted to specific phylogenetic groups, the presence or absence of peptides in a subset of species was investigated. Ten proteomes of monocots and 13 proteomes of dicots were selected as representative crops and common plant model species. Two major clades, with 0.3 and 0.11 amino acid substitution per site, were generated containing the monocots (dark blue) and the dicots (light blue) (Fig. 1A). Putative orthologues of some peptides including AtRALF23, AtPSK1, AtPIP1, and AtPSY1 were found in most monocots and dicots. For a second group, orthologues of AtPEP1, AtGRI, AtPEP3, AtPIP2, and AtPEP2 were only found in dicots. Orthologue searches for ZmPEP1 and OsIRP resulted in peptide motifs mostly specific for monocots and Zip1, NtHypSys, SlSystemin, SlCAPE1, GmPEP890, GmPEP914, and AtSCOOP12 appeared to be species-specific (Fig. 1B). Although ZmPEP1 and AtPEP1 showed certain similarities between their PR-rich N-terminus and GQHN-rich C-terminus, the motif score of the *Z. mays* orthologue of AtPEP1 was 10%, while in the reverse search using psiBLAST the orthologues of ZmPEP1 could not even be identified in *A. thaliana*. (Suppl. Fig. 3).

**Figure 1.**
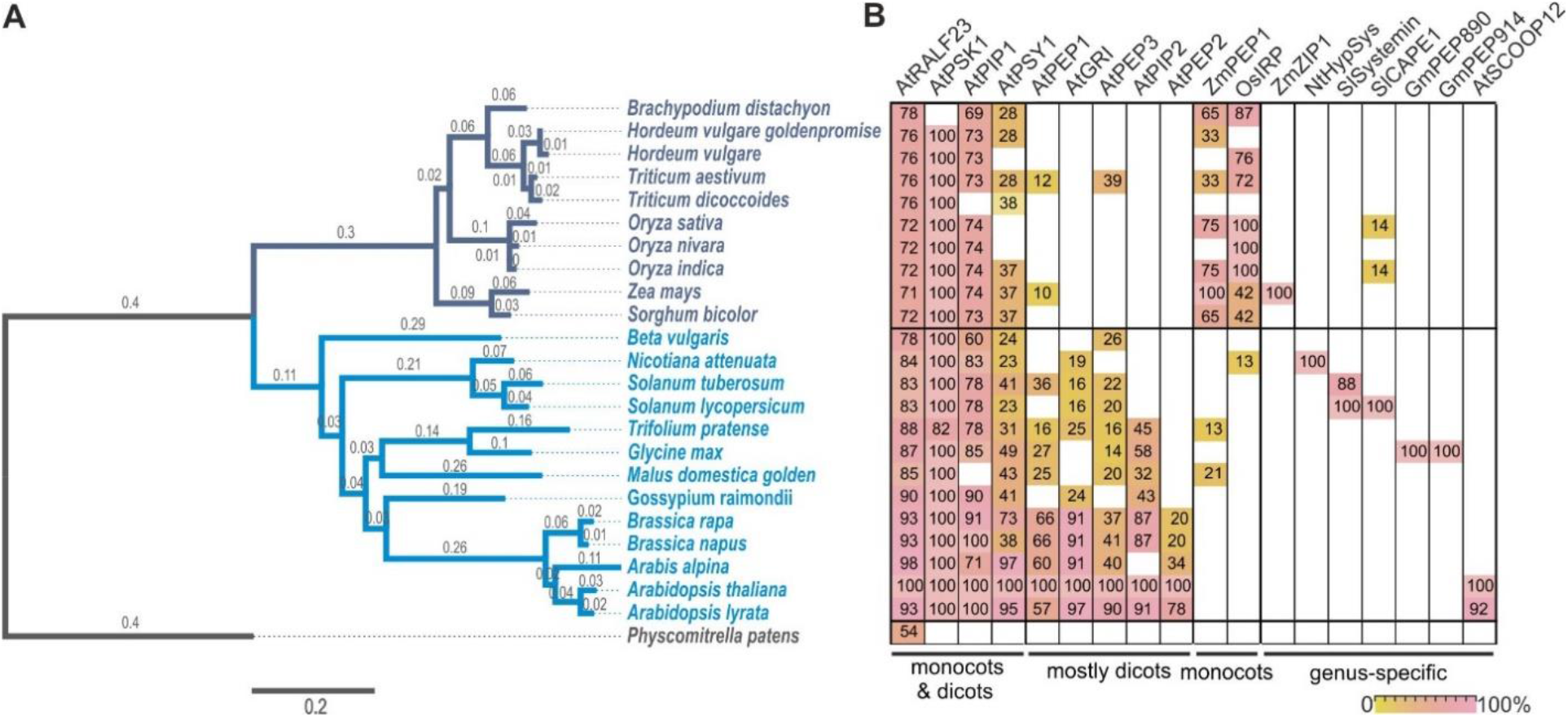
Phylogenetic analysis and distribution of phytocytokines across monocots and dicots. (**A) Phylogenetic tree of selected crops and model plants**. 23 genomes were selected as representative species of monocots (dark blue) and dicots (light blue). Phylogenetic tree was generated based on BUSCO proteins (Manni et al., 2021; Seppey et al., 2019) shared by all species and with the moss *P. patens* as outgroup. The scale bar represents the number of amino acid substitutions per site on the respective branch. **(B) Distribution of phytocytokines in selected plant species**. 18 sequence queries of known phytocytokines used for the search of orthologue sequences. Colors and values in cells indicate the motif score (0 - 100%) of the identified peptide in the respective plant species. Phytocytokines were grouped in four categories depending in which plant species they were found: monocots and dicots, mostly dicots, only monocots and genus – specific.

Altogether, six *Z. mays* phytocytokine candidates: ZmIRP, ZmPIP1, ZmPSK1, ZmPEP1, ZmRALF23 and ZmPSY1 were identified as orthologues in our analysis (Table 2). All orthologues excluding ZmPSY1 and ZmPEP1 showed a strong motif conservation across all tested species, suggesting an evolutionary diversification for these two peptides (Suppl. Fig. 4). The sequences of ZmRALF23 and ZmPEP1 have been previously described (Campbell & Turner, 2017; Huffaker et al., 2011). The identified sequence of ZmPSY1 has a motif score of 37% compared to its orthologue in *A. thaliana*, whereas the sequences for ZmIRP, ZmPIP1 and ZmPSK1 have a 42%, 74%, and 100% motif score towards the described orthologue, respectively (Fig. 1). To further characterize the biological function of the *Z. mays* phytocytokine candidates, the sequences of ZmIRP (GTDSWLESGVGMLTQLLLGAK), ZmPSK1 (AHTDYIYTQ, non-sulfated) and ZmPIP1 (RLPAGPSPKGPGH) were selected due to their higher sequence similarity to the original peptide sequence.

**Table 2:**
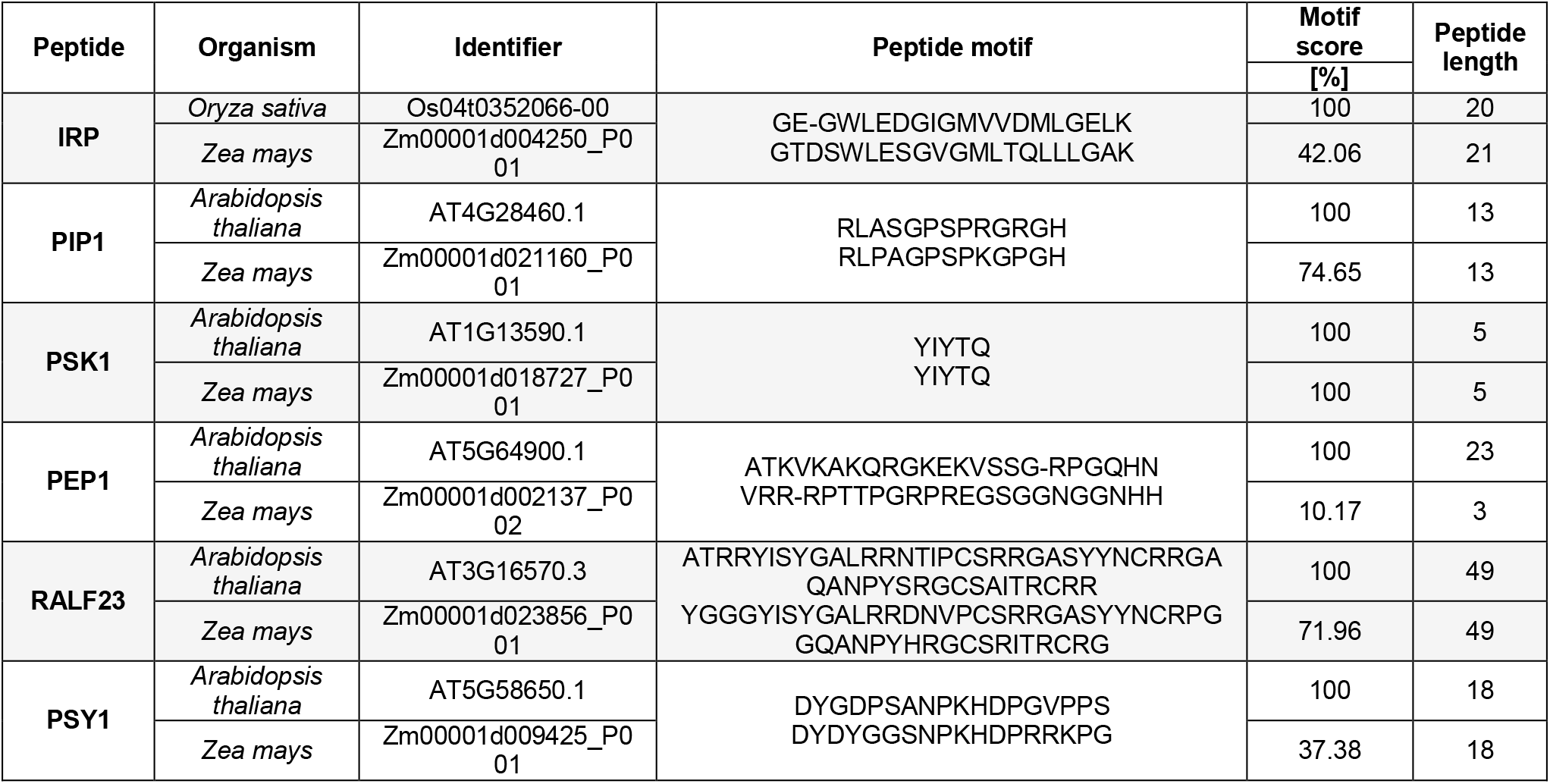
Peptide orthologues found in *Z. mays*. Six peptides identified in *Zea mays* as orthologues of known phytocytokines related to plant immunity. Peptide name, organism where the query was first identified, protein identifier, peptide sequence alignment, motif score and peptide length are shown. Hyphen represent a gap in the alignment to the query peptide.

### Maize phytocytokines trigger immune responses

To investigate a potential role of ZmIRP, ZmPSK1 and ZmPIP1 in maize, we syringe infiltrated chemically synthesized peptides into maize leaves and tested for *PR* gene expression as a readout for the activation of the immune response (Dolezal et al., 2014; Glazebrook, 2005; Ray et al., 2016; van der Linde et al., 2012). 24 hours post infiltration (hpi), leaf samples were collected and analyzed via qRT-PCR (Fig.2 A-D, Suppl. Fig. 5). As a positive control, the previously identified maize phytocytokine Zip1 (Ziemann et al., 2018) was used. As a negative control we included the maize apoplastic peptide (MAP1) (formerly described as DAMP1) which was previously shown to not trigger PR gene expression in maize (Ziemann et al., 2018). In parallel, the known MAMPs, flg22 and chitin, were used to compare the elicitation of the immune response. Both, flg22 and chitin induced a significant upregulation of all tested PR genes, confirming the elicitation of a typical MAMP-immune response in our experimental conditions. The expression of PR3, PR4, PR5 and PR10.1 was significantly up-regulated in comparison to mock after treatments with ZmIRP, ZmPSK1 and ZmPIP1 and Zip1 (Fig. 2 A, B and D; Suppl. Fig. 5). PRm6b expression was significantly induced only after ZmPSK1 and Zip1 treatments (Fig. 2C). In contrast, MAP1 did not cause a significant induction of any of the tested PR genes (Fig. 2 A - D). Together, these results show that ZmIRP, ZmPSK1 and ZmPIP1 trigger defense gene expression in maize leaves.

**Figure 2.**
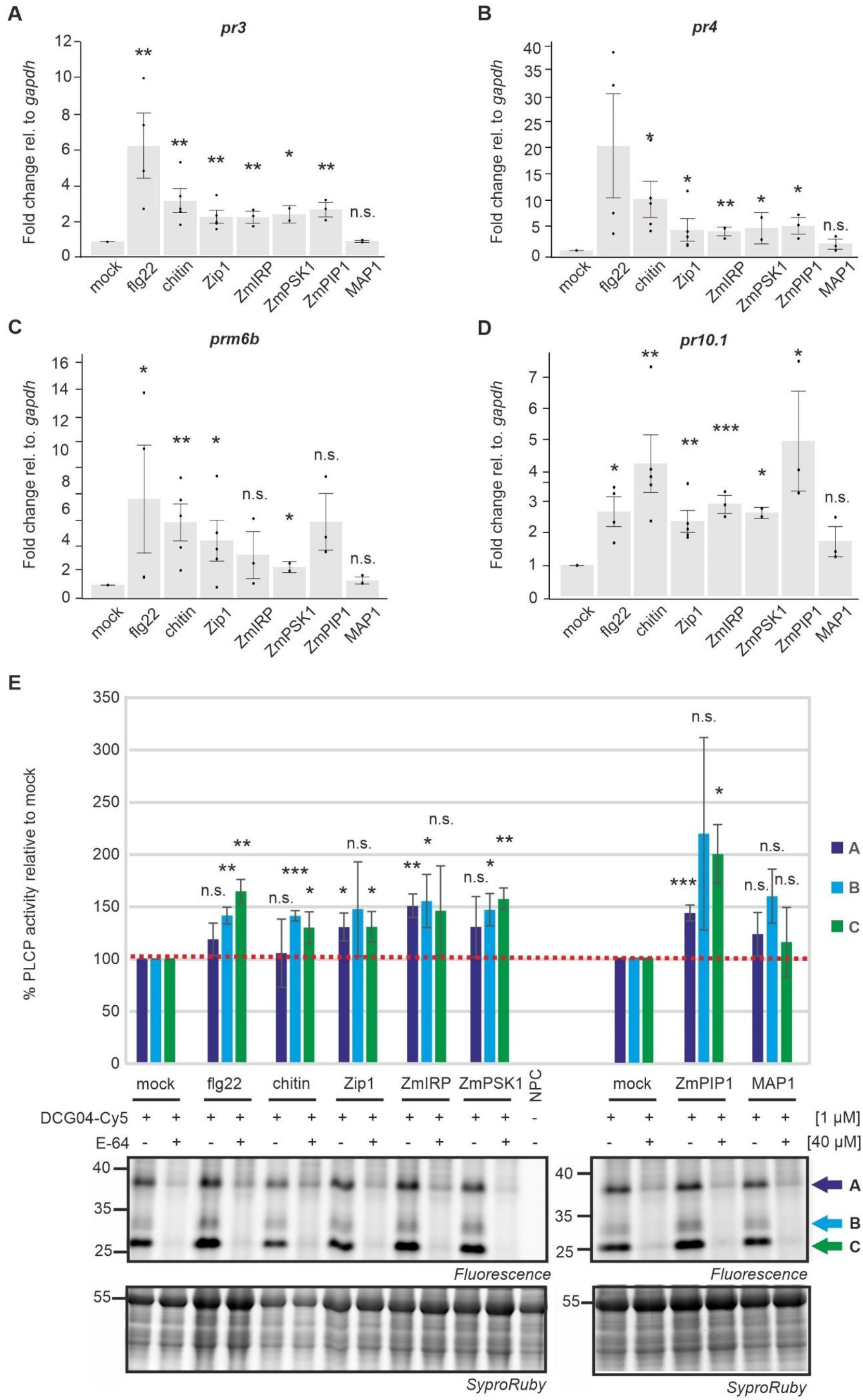
Maize phytocytokines activate defense responses. **(A-D) Maize phytocytokines induce PR gene expression**. qRT-PCR of maize leaves treated with 4 μM phytocytokines and the MAMPs flg22 and chitin. Mock and MAP1, an apoplastic derived peptide, served as negative controls. The expression of the PR genes *pr3* (Glazebrook, 2005), *pr4* (Ray et al., 2016), *prm6b* (van der Linde et al., 2012) and *pr10.1* (Dolezal et al., 2014) was analyzed 24h post infiltration. Shown are at least four independent biological replicates with three technical replicates, error bars represent SEM; p values were calculated via unpaired t-test. *p< 0.05; **p< 0.01; ***p< =0.0001; n.s. not significant. **(E) Activation of PLCPs after MAMP and phytocytokine treatment**. Activity-based protein profiling (ABPP) of maize leaves infiltrated with 4 μM synthetic peptides of the maize phytocytokines Zip1, ZmIRP, ZmPSK1 and ZmPIP1 and the MAMPs flg22 and chitin was performed 24 hpi. Mock and the MAP1 peptide were used as negative controls. Total extract of the samples was labeled at pH 6 using 1 μM DCG04-Cy5. Pre-incubation with 40 μM of E-64 shows the specificity of the PLCP signals. Samples were run on a SDS page and analyzed via fluorescent scanning using the Cy5 filter for PLCP detection. Gels were stained with SyproRuby and used as a loading control for the quantification analysis. The no-probe control (NPC) shows background signals. Gel signals (A, B and C) were quantified and values were normalized to the E-64 pre-incubation and the activity of the mock control was set to 100%. The graph shows the mean and SEM of three biological replicates whereas the gel is a representative figure.

Papain-like cysteine proteases (PLCPs) are hubs in maize immunity (Misas-Villamil et al., 2016). Therefore, we tested the activation of PLCPs after treatment with the newly identified phytocytokines in comparison to the MAMPs, flg22 and chitin. Total extracts of treated maize leaves were labeled with the activity-based probe DCG04-Cy5 for 3 hours. DCG04-Cy5 is a fluorescently-tagged derivative of E-64 that binds covalently and irreversible to the active site of PLCPs and serves to monitor the availability of active sites in complex proteomes (Greenbaum et al., 2000). Three main PLCP-specific signals can be observed between 25 and 40 KDa (Fig. 2E, bands A, B, C). Signals were quantified, values were normalized to the E-64 background (irreversible PLCP inhibitor) (Hanada et al., 1978) and the loading control and mock signals were set to 100% activity. The overall activity of PLCPs was significantly enhanced after MAMP and phytocytokine treatments, but not after MAP1 treatment. However, different PLCPs were differentially activated depending on the treatment: band A was significantly enhanced after Zip1, ZmIRP and ZmPIP1 treatments whereas band B showed stronger signals for all treatments except for ZmPIP1 and Zip1. Signals for band C were stronger after flg22 and chitin treatment as well as for all phytocytokines except for ZmIRP (Fig. 2E). Thus, MAMPs and phytocytokine treatments activate distinct PLCPs, which in turn might trigger differential immune responses. Overall, these data indicate that the newly identified peptides ZmIRP, ZmPSK1 and ZmPIP1 elicit maize defense responses and might function as phytocytokines in maize immunity.

### Phytocytokines induce the up-regulation of phytohormone - related genes

It has been previously shown that phytocytokines are involved in phytohormonal immune signaling (Hou et al., 2021). We therefore tested, which hormonal pathways become activated in response to our set of peptides. Three and 24 hours after treatment, leaf samples were collected and gene expression was analyzed via RT-qPCR. As marker for the SA signaling pathway the genes *atfp4*, *pox12* (Hemetsberger et al., 2012), *acd6/ank23* (Zhang et al., 2019), *pal1* (Wang et al., 2020), and *wrky65* (Huo et al., 2021) were tested. For the JA pathway the genes *myc7e* (Engelberth et al., 2012), *igl1* (Frey et al., 2004), *mpi1* (Borrego & Kolomiets, 2016) and *cc9* (van der Linde et al., 2012) were analysed. The MAMPs, chitin and flg22 activated SA--related marker genes (Fig. 3A - F). At 3 hpi, chitin caused a downregulation of *igl1*, which is involved in dimboa synthesis and associated to the JA pathway (Fig. 3E). flg22 triggered an upregulation of *wrky65*, a transcription factor associated to the SA signaling pathway (Fig. 3B). At 24 hpi the SA markers *pal1* and *ank23* were upregulated in response to chitin, whereas the *pal1* and *pox12* induced an upregulation after flg22 treatment (Fig. 3A,D,F). In addition, the newly identified maize phytocytokines also triggered the expression of hormonal marker genes. ZmIRP significantly triggered the upregulation of the SA markers *ank23* at 3 hpi and 24 hpi as well as of *pal1* and *pox12* at 24 hpi (Fig. 3A,D,F). The JA marker *myc7e* was upregulated after ZmIRP treatment at 24 hpi (Fig. 3C). Besides, ZmIRP treatment caused downregulation of *igl1* at 3 hpi (Fig. 3E), correlating to an activation of SA defense responses. ZmPSK1 treatment caused an upregulation of the three SA marker genes *ank23, wrky65* and *pal1* at 24 hpi, although at 3 hpi, *pox12* and *igl1* expression was downregulated (Fig. 3A-F). ZmPIP1 did only upregulate the expression of *wrky65* at 24 hpi (Fig. 3B). Zip1 upregulated the SA biosynthesis gene *pal1* at 24 hpi, which is in line to the previous report that Zip1 induces SA responses (Ziemann et al., 2018). Gene expression analysis of marker genes for SA- and JA-related immune pathways suggest that both, MAMPs and phytocytokines activate immune components of hormonal pathways, but to a different strength and often antagonistic responses.

**Figure 3.**
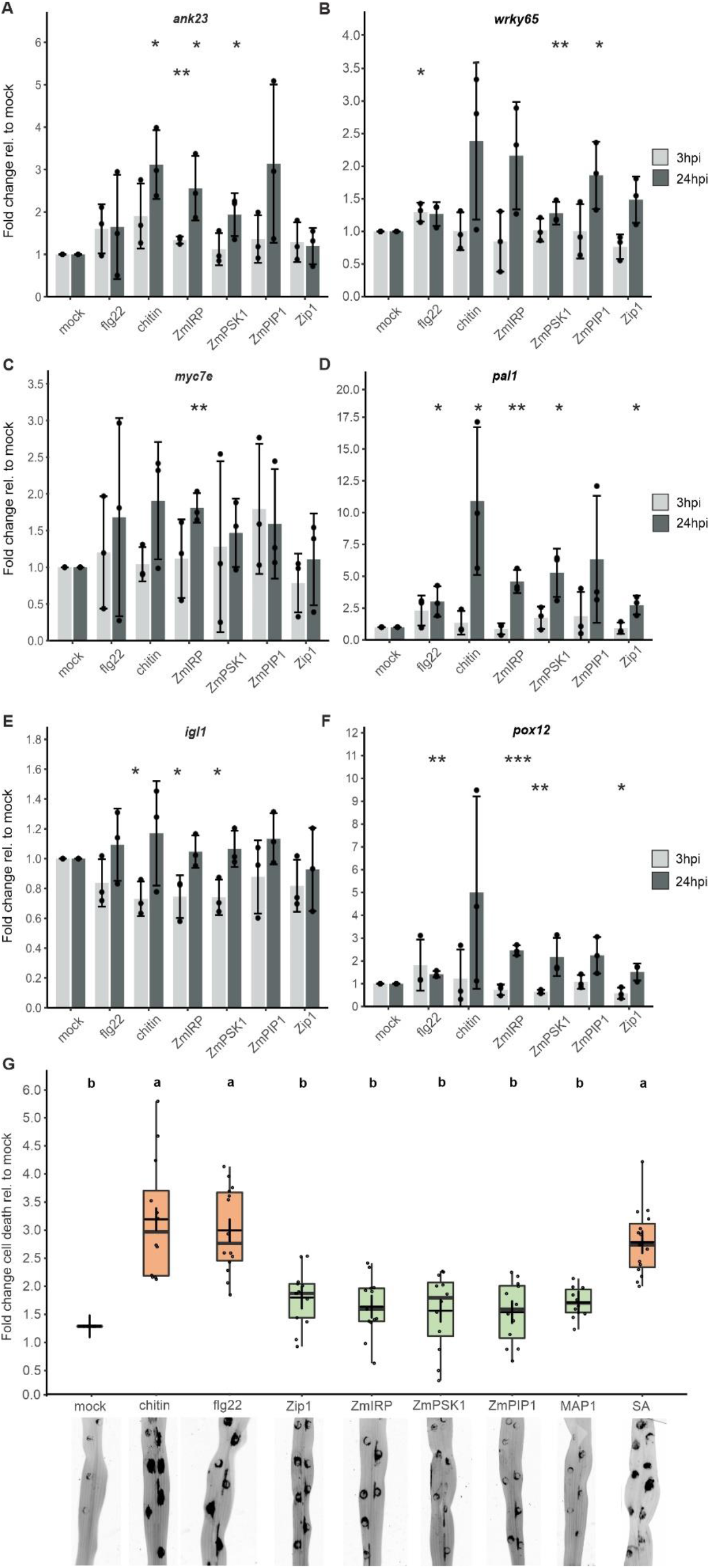
Maize phytocytokines trigger phytohormonal pathways but do not promote cell death. **(A - F) Gene expression analysis shows differential regulation of hormonal pathways**. The expression of the marker genes *ank23* (Zhang et al., 2019) (A), *wrky65* (Huo et al., 2021) (B), *myc7e* (Engelberth et al., 2012) (C), *pal1* (P. Wang et al., 2020) (D), *igl1* (Frey et al., 2004) (E) and *pox12* (Hemetsberger et al., 2012) (F) was tested after maize leaves were treated with the MAMPs, flg22 and chitin, the phytocytokines ZmIRP, ZmPSK1, ZmPIP1 and Zip1 or the mock solution at 3 and 24 hpi. Gene expression of at least three biological replicates was normalized to *gapdh* and the fold change expression relative to mock was analysed. Shown are at least three independent biological replicates with two technical replicates, error bars represent SD; p values were calculated via unpaired t-test. *p< 0.05; **p< 0.01; ***p< =0.0001. **(H) Maize phytocytokines, in contrast to MAMPs and SA, do not induce cell death in co-incidence with wounding**. Cell death quantification at 24 hpi of maize leaves treated with 4 μM flg22, 4 mg / ml chitin or 4 μM of the synthetic peptides Zip1, ZmIRP, ZmPSK1 and ZmPIP1. Mock and MAP1 served as negative controls. Additionally, 2 μM salicylic acid (SA) was tested. Shown are representative pictures imaged 24 hpi with a blue epi-illumination source. Black signals represent cell death. The results of at least three independent biological replicates each with six technical replicates were plotted. Data analyses were performed using an ANOVA. Letters above indicate significant differences between samples (α = 0.05, Tukey test).

### Phytocytokines do not induce cell death in co-incidence with damage

It was recently shown that simultaneous perception of MAMPs and cell damage triggers a strong localized immune response in Arabidopsis roots (Zhou et al., 2020). We speculated that leaves would also similarly react to this combination of triggers and a strong cell death response might occur. At the same time, we asked, if not only MAMPs, but also phytocytokines could result in a similar output. A combination of damage and MAMPs, or damage and phytocytokines was tested in maize leaves syringe infiltrated with the MAMPs, flg22 and chitin, and with the maize phytocytokines Zip1, ZmIRP, ZmPSK1 and ZmPIP1 and the control peptide MAP1. The syringe infiltration caused damage at the infiltrated site from its own, which can be observed as a circular lesion in the form of a syringe without a needle (Fig. 3G). Autofluorescence with a GFP filter was used as a measurement for the quantification of cell death 24h post infiltration. SA was included as additional treatment, since a variety of phytocytokines and MAMPs trigger immunity through the SA defense pathway (Hou et al., 2021). At 24 hours after treatment, flg22 and chitin caused strong cell death at the infiltration site compared to mock (Fig 3G). Also, the treatment with SA induced a similar spread of cell death as the MAMP treatments (Fig. 3G). In contrast, none of the tested phytocytokines, nor the control peptide MAP1, caused a significant induction of cell death in comparison to the mock sample (Fig. 3G). These findings demonstrate that the combination of damage and MAMPs (or SA) trigger a cell death response in maize leaves in contrast to the phytocytokines, which did not trigger cell death. Thus, we conclude that although both MAMPs and phytocytokines activate overlapping defense responses related to phytohormonal pathways, the signaling output with regard to cell death differs between both triggers.

### Phytocytokines alter virulence of biotrophic and necrotrophic pathogens

To gain further insights into the roles of the maize phytocytokines ZmIRP, ZmPSK1 and ZmPIP1 in plant immunity, we conducted infection assays with the necrotrophic fungal pathogen *B. cinerea*. To this end, *B. cinerea* conidia were applied onto the infiltrated leaf tissues 24 h after infiltration with the tested peptides. Subsequently, lesion sizes were measured at 48 hours after fungal inoculation. Treatment with the MAMPs, flg22 and chitin, resulted in significantly reduced lesion size compared to mock (Fig. 4A) due to the induction of defense responses. Similarly, ZmPSK1 treatment significantly reduced the lesion size in a similar manner as the chitin treatment, suggesting that ZmPSK1 enhances resistance against *B. cinerea*. Contrary, the phytocytokines ZmIRP and Zip1 caused a significant increase in *B. cinerea* lesion size, indicating enhanced susceptibility of the maize leaves upon treatment with these peptides. ZmPIP1 did not cause a statistically significant effect compared to mock control or the MAP1 treatment (Fig. 4A). This experiment revealed that both, MAMPs and phytocytokines alter plant susceptibility towards the necrotrophic pathogen *B. cinerea*. While both tested MAMPs, as well as ZmPSK1 appeared to cause an increased resistance towards *B. cinerea*, the phytocytokines Zip1 and ZmIRP promoted infection of the necrotrophic pathogen.

**Figure 4.**
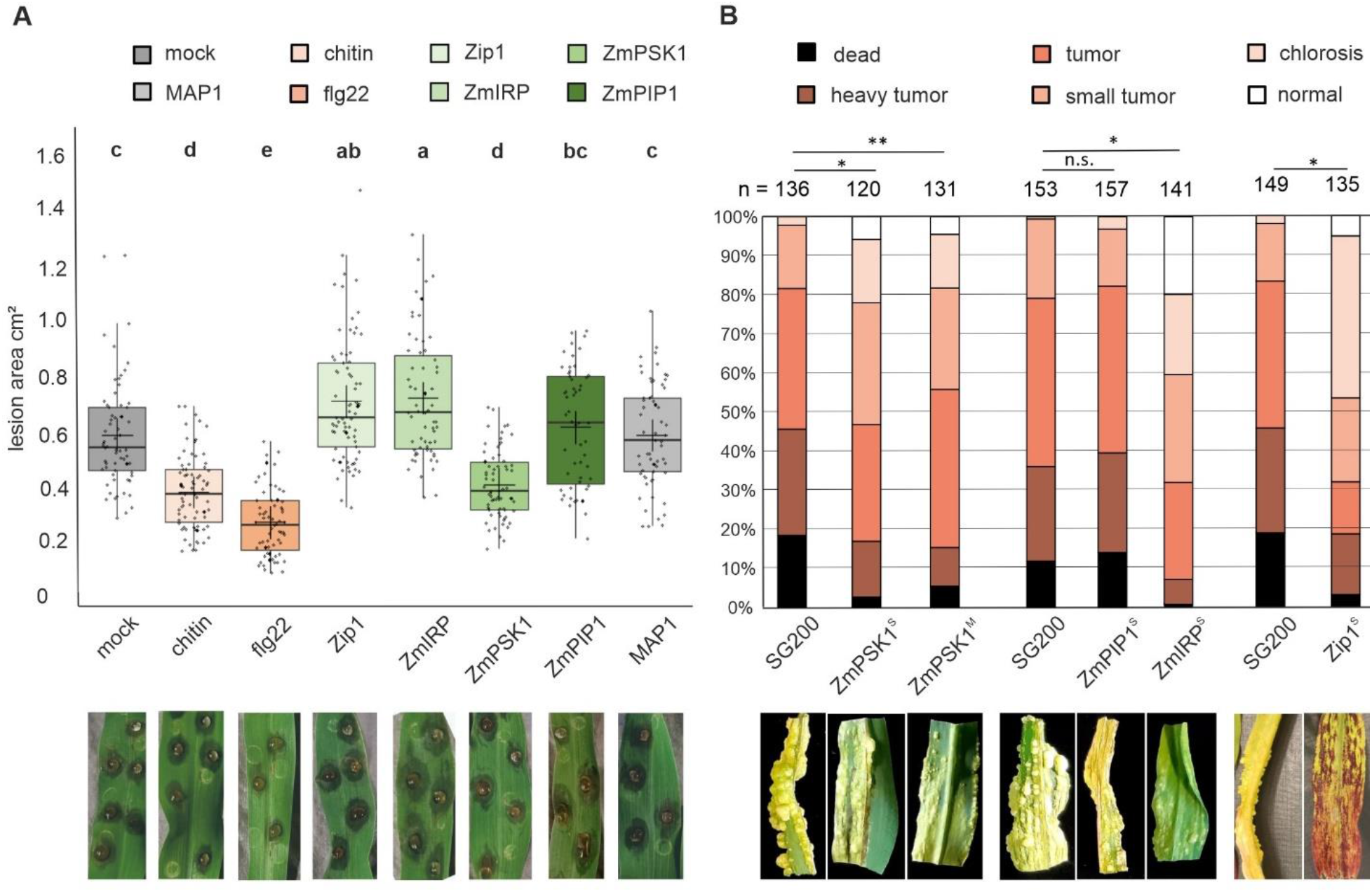
Maize phytocytokines influence virulence of necrotrophic and biotrophic fungal pathogens. **(A) Maize phytocytokines alter lesion size upon *B. cinerea* infection**. Lesions of maize leaves treated with 4 μM phytocytokines, MAMPs and the control peptide AMP1 were quantified 48 hpi. Shown are representative pictures of the infection at 48 hpi. This experiment was performed with at least three independent biological replicates each with six technical replicates. Statistical analysis was performed using an ANOVA. Letters above indicate significant differences between samples (α = 0.05, Tukey test) **(B) Presence of maize phytocytokines affect *U. maydis* virulence**. Seven days-old maize seedlings were infected with the solo-pathogenic strain SG200 and transgenic SG200 expressing the phytocytokines ZmPSK1, ZmPIP1, ZmIRP and Zip1 under the control of the *pit2* promotor. Single (s) and multiple (m) integrations into the SG200 genome were tested. Disease symptoms were quantified and representative pictures of the infection were taken at 12 dpi. Statistical analyses were performed with a T-test (*p< 0.05; **p< 0.01; ***p< =0.0001). n = number of tested plants. Shown are the results of at least three independent biological replicates.

To test the responses of a biotrophic pathogen to the maize phytocytokines we made use of the “trojan horse” approach in which the fungus *Ustilago maydis* is deployed to deliver plant peptides into the leaf apoplast *in-vivo* (van der Linde et al., 2018; Ziemann et al., 2018). *U. maydis* infects primordia of all aerial organs of maize and virulence can be evaluated by the induction of chlorosis, anthocyanin and tumor formation (Schilling et al., 2014; Skibbe et al., 2010). We generated recombinant *U. maydis* strains secreting ZmPSK1, ZmPIP1 and ZmIRP phytocytokines during biotrophic infection. In addition, a previously generated *U. maydis* strain that delivers the Zip1 peptide was used as a positive control. Expression constructs for the plant peptides were integrated as a single copy into the *U. maydis* genome. In addition, we selected *U. maydis* transformants that contained multiple integrations of ZmPSK1, which allowed to test for an eventual dose effect of the delivered phytocytokine. Maize seedlings were infected with the recombinant fungal strains and the resulting disease symptoms were quantified at 12 dpi. Secretion of Zip1 resulted in a reduction of *U. maydis* disease symptoms, confirming previous results (Fig. 4B) (Ziemann et al., 2018). Similarly, ZmPSK1 had a negative impact on *U. maydis* infection, and the expression of ZmIRP resulted in even stronger reduction of *U. maydis* virulence (Fig. 4B). In contrast, ZmPIP1 seemed to enhance *U. maydis* virulence, although not statistically significant in comparison to the SG200 infection (Fig. 4B). Thus, ZmPSK1 and ZmIRP significantly interfere with *U. maydis* infection restricting disease symptoms. Together, we observed that maize phytocytokines modulate infection of both, necrotrophic and biotrophic pathogens in a differential manner. ZmIRP and Zip1 enhance resistance to the biotrophic pathogen, but increase susceptibility to the necrotroph correlated to an activation of SA-related defense responses. In contrast, ZmPSK1 reduces virulence of both, biotrophic and necrotrophic pathogens, whereas ZmPIP1 did not interfere with *B. cinerea* infection nor promoted tumorigenesis caused by *U. maydis*.

## Discussion

In this study we have used a bioinformatics pipeline to identify six maize orthologues of phytocytokines from other plant species, three of which were confirmed as biologically active signaling peptides in maize: ZmIRP, ZmPIP1 and ZmPSK1. Similar to the MAMPs, flg22 and chitin, these peptides trigger maize immune responses such as PR-gene expression and PLCP activation. ZmIRP and ZmPSK1 differentially activate hormonal pathways thus playing an important role in the plant immune response against biotrophic and necrotrophic pathogens. ZmIRP, similarly to Zip1, has been shown to activate the SA pathway (Wang et al., 2020). Consequently, both peptides confer resistance to the biotrophic pathogen *U. maydis*, but enhance susceptibility towards the necrothoph *B. cinerea*, reflecting the antagonistic SA-JA dependency of plant resistance towards biotrophic vs necrotrophic pathogens (Glazebrook, 2005). Thus, in our experiments ZmIRP and Zip1 appear to have a similar impact on the maize immune response.

Perception of MAMPs by membrane localized PRR receptors leads to activation of signaling pathways related to PTI-responses (DeFalco & Zipfel, 2021; Hou et al., 2021). Some phytocytokines trigger PTI-related responses, suggesting overlapping components being involved in perception and upstream signaling. For instance, Arabidopsis AtPIP1 (PAMP-induced secreted peptide 1) induces MPK6 and MPK3 activation, ROS production, callose deposition and enhanced *frk*1, *wrky53* and *wrky33* gene expression as well as other SA-related components (Hou et al., 2014; Hou, Shen, et al., 2019). Expression of *propip1* is upregulated by flg22, *Pseudomonas syringae* DC3000 and *Fusarium oxysporum* infection, which results in enhanced resistance against these pathogens. AtPIP1 has been therefore been proposed as an amplifier of flg22 responses (Hou et al., 2014). In maize, we found that ZmPIP1 triggers defense responses such as *PR*-gene expression and PLCP activation, but it does neither lead to a significant upregulation of SA- or JA-responsive genes, nor to an altered resistance towards the tested plant pathogens.

Differences in phytocytokine responses could also indicate that their perception and function differ between plant species, i.e., between monocots and dicots. Indeed, PEPs were not recognized by species outside of their plant family of origin, likely due to a divergence of the PEP sequences and adaptation of the PEPR receptors (Lori et al., 2015). In contrast to maize, where PEPs activate distinct signaling phytohormonal pathways such as JA, ETH, volatile terpenes and indoles (Poretsky et al., 2020), in Arabidopsis the PEPs have largely redundant activities (Bartels et al., 2013; Huffaker et al., 2006). This indicates that, although PEPs are conserved as plant immune components, there are regulatory differences among species (Poretsky et al., 2020). In maize, Zip1 responses are distinct from what is described for phytocytokines in Arabidopsis. In particular, Zip1 does not trigger any of the canonical PTI responses, but instead activates transcriptional reprogramming and PLCP activation largely overlapping with SA responses (Ziemann et al., 2018).

### MAMP and phytocytokine responses diverge in cell death

In the course of this study we observed that maize phytocytokines activate plant defense genes similar to MAMPs, however, in co-incidence with wounding, the immune responses bifurcate: While the combination of wounding and MAMPs resulted in cell death, this was not the case for the phytocytokines. In Arabidopsis roots it has been shown that damage of small cell clusters strongly upregulates PRR expression (Zhou et al., 2020). This leads to a localized immune response of otherwise non-responsive cells to beneficial bacteria, indicating that damage “switches-on” local immune responses, generating more PRRs capable of sensing MAMPs. It has been proposed that the levels of signals perceived might contribute to the diversification and specificity of the immune response (Hou et al., 2021), but this also might suggest that the number of available PRRs could be the regulating factor. Indeed, perception of the tyrosine sulfated peptides ROOT MERISTEM GROWTH FACTOR (RGF)/GOLVENs (AtGLVs) in Arabidopsis promotes the stability and abundance of the FLS2 and EFR receptors (Stegmann et al., 2022) and external application of AtPIP1 also induces transcriptional upregulation of PRRs (Hou et al., 2014; Rhodes et al., 2021). After phytocytokine perception PRRs could be stabilized in a spatio-temporal manner making surrounded cells more responsive for the invaders and ready to futher amplify the immune response. Besides, phytocytokines could also play an important role in the healing process of the cell (Vega-Muñoz et al., 2020). A(nother) possible scenario is that the faith of the cell is decided downstream of perception. This hypothesis is supported by a recent finding in Arabidopsis showing that both, ETI- and PTI-defense pathways are activated downstream of BAK1 receptor recognition (Schulze et al., 2022). With the available information, one could hypothesize that phytocytokines are used by the plant as alert signals for cell-to-cell communication to amplify immune responses in by-stander cells as a survival mechanism (Fig. 5). Indeed, only few reports describe the existence of “death” peptides in plants. One example is the kiss-of-death (AtKOD) peptide from Arabidopsis which is upregulated by biotic and abiotic stresses and its expression causes cell death in leaves and seedlings (Blanvillain et al., 2011). Another example in Arabidopsis is the GRIM REAPER (AtGRI) peptide which has been shown to be activated by the MC9 releasing a 11 amino acid peptide which is then recognized by the receptor-like kinase PRK5 inducing ROS dependent cell death (Wrzaczek et al., 2015). Future experiments will aim to unravel the cellular components that determine the initiation of immune-regulated cell death, or rather promote a phytocytokine-driven pro-life decision in immune-activated cells.

**Figure 5.**
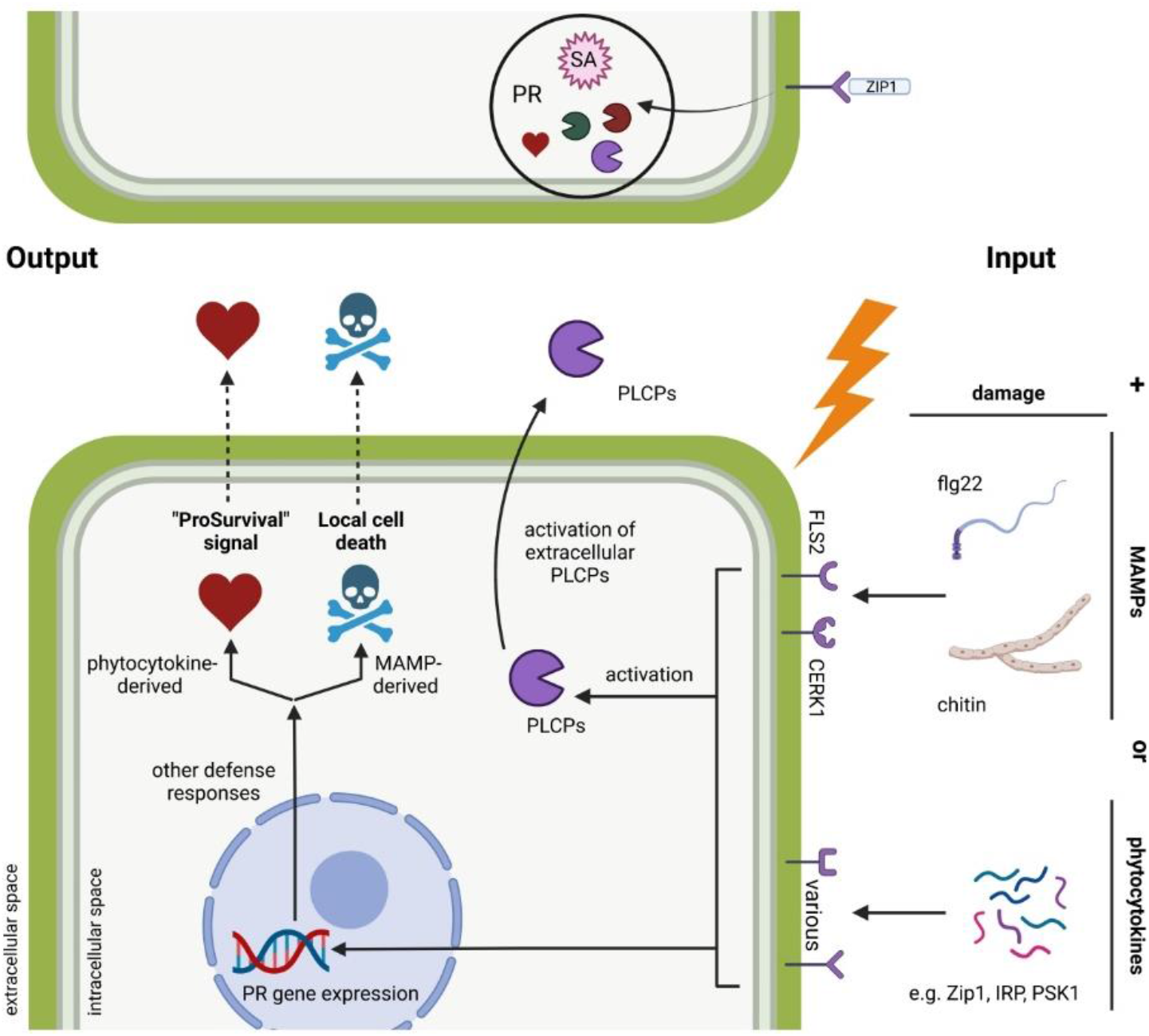
Model of phytocytokine immune activation. MAMPs and phytocytokines are recognized by surface receptors triggering immune responses such as induction of PR gene expression and PLCP activation. In co-incidence with wounding (input), recognition of MAMPs and phytocytokines activate differential pathways in- or out-side the surrounding cells leading to a pro-survival signal (output) triggered by the phytocytokines in contrast to a local cell death signal (output) induced by MAMPs.

## Material and Methods

### *In silico* identification of putative propeptides and peptide hormones

A BLAST database consisting of predicted proteomes from 78 randomly selected plants including 31 monocots, 43 dicots and 4 from other phyla was created (Suppl. Table 1). The proteomes were obtained from the plants.ensemble.org database. Eighteen previously published propeptides (Table 1) were used in all BLASTs as queries in the NCBI BLAST+ application (Camacho et al., 2009). psiBLAST searches (Altschul et al., 1997; Schäffer et al., 2001) of the queries in the database were performed with five iterations and an e-value threshold of 0.05 to detect distant relationships between proteins. A parallel BLASTp (Altschul et al., 1990) with the same settings was performed.

After the BLASTs, the results for eighteen BLAST searches were combined, and unique identifiers were assigned to each hit linking it to the respective BLAST search. Redundant hits for every proteome were removed after sorting all hits by their bit score and selecting only the hit with the highest score. Next, low-quality hits were removed from BLASTp and psiBLAST results: Hits with less than 10% protein identities and query coverage of less than 25% in the psiBLAST and BLASTp were removed, respectively. To focus on only the hits of each organism with the highest homology to the query, only the best psiBLAST hits per organism were selected after hits were sorted by their bit scores.

Due to variance in the sequence length of all hits, hits whose amino acid sequence length was 1.5-times longer or 1.5-times shorter than the longest or shortest hit within the group of hits with more than 75% identity, respectively, were removed. In the last step, hits were filtered according to their similarity to the respective peptide hormone of each query. All hits for each queried propeptide were aligned with the ClustalOmega (Goujon et al., 2010) application, and the region containing the peptide hormone was analyzed and scored. The similarity of each alignment region containing the peptide hormone was scored based on a BLOSUM62 matrix (Henikoff & Henikoff, 1992) and low gap costs (gap: −8, extension: −1) using the biopython package (Cock et al., 2009) (Equation 2):

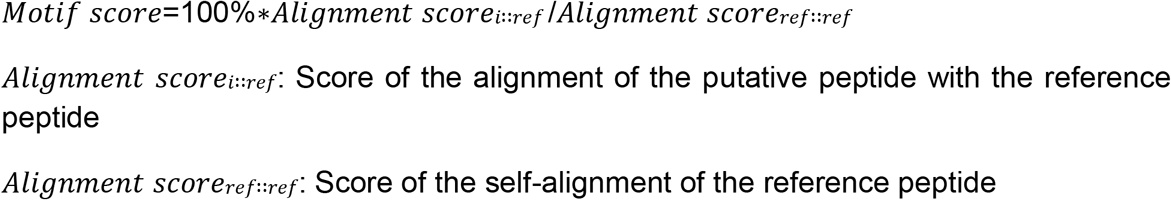

To account for the differing length and AA compositions of the peptide hormones, the final so-called motif score was calculated relative to the scoring of the self-aligned peptide hormone. Hits with motif scores lower than 10 % were removed. The process of alignment, scoring, and filtering was performed three times. The phylogenetic tree was created as following based on the proteome of ten monocots and 13 dicots which were selected as representative crops and common plant model species: 140 complete proteins present in all proteomes of the selected organisms either as one or multiple copies were selected using the BUSCO software (Manni et al., 2021; Seppey et al., 2019) in its standard settings. If multiple copies were present, only the best hit was used to create the phylogenetic tree. The orthologs of each BUSCO protein were aligned via ClustalOmega. The alignments of all proteins were concatenated. A phylogenetic tree of the concatenated multiple sequence alignments was calculated via RAxML (Stamatakis, 2014) using the “PROTCATBLOSUM62” substitution model. The phylogenetic tree was visualized with FigTree (Andrew Rambaut, 2018) using *Physomitrella patens* as the outgroup. Plots of results regarding peptide hormones were created with the matplotlib (Hunter, 2007) and the logomaker (Tareen & Kinney, 2019) packages in python. The venn diagram was created with the “ggvenn” package in R.

All scripts are deposited on github.com: https://github.com/dmoser1/phytocytokine.git

### Plant growth conditions

Grains of maize (*Zea mays*, cv. Golden Bantam) were placed in organic growth soil and grown for seven to eight days in a plant growth chamber (15 h photoperiod at 28°C, 1h twilight morning and evening and 7 h darkness at 22°C).

### Plant treatments

#### Chitin

50 mg Shrimp shell chitin (Sigma Aldrich C9752) was grinded in 2.5 mL ddH_2_O with mortar and pestle. Residues were flushed with additional 2.5 ml ddH_2_O. The mortar was placed in a microwave for 60 seconds. The chitin solution was transferred to a 15 ml falcon tube and filled up to 5 ml with ddH_2_O to compensate evaporation. The solution was sonicated using a microtip (Sonoplus KE76) for three minutes using pulse and 70% amplitude. The sonicate was centrifuged for five minutes at 2500 g and the supernatant (10 mg / ml) was transferred to a new tube. For the treatments the supernatant was diluted to a final concentration of 4 mg / ml using ddH_2_O.

#### flg22

1 mM flg22 was diluted to a final concentration of 4 μM in ddH_2_O. ZmIRP (GTDSWLESGVGMLTQLLLGAK),MAP1 (IFDDGGFGEVHADPIKVER),ZmPSK1 (AHTDYIYTQ, non-sulfated),ZmPIP1 (RLPAGPSPKGPGH),Zip1 (EGESELKLATQGASVRR): Lyophilized peptide powder was diluted to stock concentrations of 0.5 mg / ml. The final concentration to treat the plants was 4 μM each. ZmPSK1 peptide is not sulfated.

The second leaf of eight days-old maize seedlings was infiltrated using 1 ml needleless syringe.

### PR gene expression analyses

Treated leaf samples were taken 24 hours post infiltration and grinded with liquid nitrogen. RNA was extracted using TRIzol® following the protocol of Ambion Life technologies. RNA was treated by DNase using “TURBO DNA-free” reagents following the corresponding protocols. 2 μg RNA was transcribed to cDNA using “RevertAid H minus First strand cDNA synthesis Kit” Thermo Scientific #K1632. Oligo (dt)_18_ primer were used for the synthesis. The obtained cDNA was diluted to 1:100 using Nuclease-free water. The diluted cDNA was used for qRT-PCR analyses with gene specifc primers (Suppl. Table 2). We used IQSYBRGreen (PROMEGA) reagent for detection. GAPDH was used as housekeeping gene in maize experiments. *Pr3*, *pr4*, *pr5*, *prm6b* and *pr10.1* were included in the analysis as defense marker genes. *Igl1*, *ank23*, *myc7E*, *wrky65*, *cc9*, *pox12* and *pal1* were included in the analysis as SA or JA marker genes.

### PLCP activity assay – Activity-based protein profiling (ABPP)

Treated leaf samples were taken 24 hours post infiltration and equal amounts of leaf powder was dissolved in 50 mM NaOAc buffer (pH=6) using VIBRAX shaker system (IKA) for 3 x 1 min. Samples were centrifuged 17.000g 20 min at 4°C and the supernatant was transferred to new tubes and kept on ice. 50 μL labelling reactions were prepared and 48.5 μL of protein extract was used together with either 1 μL DMSO or E64 (PLCP inhibitor) (Hanada et al., 1978) and 0.5 μL 1 M DTT. After 30 min pre-incubation 1 μL of 100 μM DCG04-Cy5 probe (obtained from Hermen Overkleeft) was added (Greenbaum et al., 2000). The samples were labelled for three hours at room temperature in the dark. The labelling process was stopped by the addition of 15 μL Laemmli buffer and heating for ten minutes at 98°C. 15 μL of sample were loaded on 15% PAA gels and ran for 70 min at 200 V in the dark. The ABPP signals were analyzed immediately after the run using a ChemiDoc (BIORAD) system with a Cy5 suitable setting. Afterwards the gels were incubated in SyproRuby fixation solution (50% MeOH, 7% acetic acid) for 30 min and incubated overnight in SyproRuby staining solution (Thermo Fisher Scientific - S12000). The following day the gels were washed applying SyproRuby washing solution (10% MeOH, 7% acetic acid) for 20 min. The signals were detected afterwards using the SyproRuby setting from the ChemiDoc. ABPP and SyproRuby signals were quantified using the ImageLAB software. A conserved protein band at 55 kDA was used to determine the relative protein concentration for each sample. The calculated correction factor was used to compare and analyze PLCPs signals for A, B, C between the mock sample and samples of treatments. Mock PLCP values have been set to 100% and accordingly the samples of the respective treatments are depicted in percent of PLCP activity.

### *Botrytis* cultivation and infection assays

*B. cinerea* strains were cultured on agar containing malt extract (ME) medium. Conidia were scraped off the 10 days-old plates with 10 ml water using a glass spatula, filtered through Miracloth (VWR), and counted using a hemocytometer. Infection tests were performed with detached 2^nd^ leaves of nine days-old maize (*Zea mays*, cv. Golden Bantam) seedlings. Prior infection, the seedlings were treated with MAMPs or phytocytokines as described previously and leaves were detached 24h post infiltration. Leaf inoculations were performed using 10 μL droplets with 2*10^5^ conidia per mL in GB5 minimal medium (GB5: 3.05g/L GB5, 10mM KH2PO4, pH 5.5). Multiple droplets were placed on individual leaves and considered as technical replicates. For one biological replicate at least three leaves were used for a single treatment. The lesion area was quantified 48 hpi using ImageJ software.

### Generation of Ustilago-phytocytokine overexpression lines

Corresponding sequences of the peptides Zip1, ZmIRP, ZmPSK1 and ZmPIP1 were cloned into the p123 vector system using the Gibson assembly method (Gibson et al., 2009). The vector harbors a constitutive promoter (pro^pit2^) for overexpression, a terminator (Tnos) and a resistance marker cassette (carboxin (cbx)). The assembled vectors were transformed into TOP10 *E. coli* cells by heat shock for plasmid multiplication. Purified plasmids were digested by SspI restriction enzyme for linearization to allow the homologous recombination into the *ip* locus. 50 μL protoplasts of *Ustilago maydis* (SG200; solo pathogenic mutant) were transfected by adding 5 μg of linearized plasmid, 1 μL Heparin and 0.5 mL STC / 40%PEG (sterile) incubating for 15 min on ice. The samples were plated on Regeneration agar (Reg-agar) - poured as two-layer system (without (top) and with cbx (bottom). Plates were incubated at 28°C for 4-5 days. Colonies were picked and singled out onto Potato-Dextrose-agar (PDA) plates for two days at 28°C. 2 mL overnight cultures of singled out colonies were used for DNA extraction (Hoffman & Winston, 1987). The extracted DNA was digested by HindIII, separated on 0.9% agarose gel (100 V, 2h) and blotted on nylon membrane for Southern Blot analysis. The detection of cbx-containing DNA fragments was performed using digoxigenin (DIG) antibody targeting the *ip* locus via a cbx probe. Successfully confirmed strains were used for infections.

### Ustilago cultivation and infection assays

SG200, SG200_Zip1, SG200_ZmIRP, SG200_ZmPSK1 and SG200_ZmPIP1 were grown to an OD_600_ of 0.8 in 55 mL YEPS_light_. The cultures were pelleted at 3500 rpm for seven minutes and the supernatant discarded. The pellets were washed with 20 mL ddH_2_O. At last, the pellets were resuspended in the calculated volume of water to reach an OD_600_ of 1.0. 10 μL droplets of each culture were placed on charcoal plates to check for filamentation (pathogenicity). 7 days-old maize seedlings were infected using a 1 mL syringe with needle injecting *Ustilago* suspension into the stem of the seedlings. The plants phenotype was investigated 12 dpi using a disease rating chart for *Ustilago maydis* comprising healthy plants, chlorosis, small tumors (<2 mm), normal tumors (2 to 10 mm diameter), heavy tumors (>10 mm or stunted growth) and dead plants. The plants were categorized based on the named phenotypical symptoms and compared to SG200 for the analyses.

### Cell death assays

MAMP- and phytocytokine-treated leaf samples were harvested 24 hpi and scanned for autofluorescence using the ChemiDoc (BIORAD) autofluorescence settings (530 / 280nm filter). The images were analyzed by ImageLAB software. Non-infiltrated areas of leaves were selected as background signal and compared to the overall signal intensity of the whole leaf surface. The signal intensity of mock (water-treated) leaves was plotted against MAMP- and phytocytokine-treated leaf samples. Mock has been set to 1 and was compared to signals of the remaining samples, depicting the fold changes relative to mock.

## Acknowledgments

We would like to thank Prof. Herman Overkleeft for providing the DCG04-Cy5 probe and Janina Werner for helping with *Botrytis* infection assays. We are also grateful to Ute Meyer for technical support. We acknowledge support from the Cluster of Excellence on Plant Sciences (CEPLAS) funded by the Deutsche Forschungsgemeinschaft (DFG, German Research Foundation) under Germany’s Excellence Strategy – EXC 2048/1 – project ID: 390686111. Our research is funded by the DFG through SFB1403 (project number 414786233) and DFG project DO 1421/5-2. Jasper R.L. Depotter was supported by the Research Fellowship Programme for Postdoctoral Researchers of the Alexander von Humboldt Foundation.

## Author contributions

MK, DM, JD, GD and JCMV designed the study. DM and JD performed the bioinformatics analysis. MK, DM and JL performed the experiments and the data analysis together with JD, GD and JCMV. MK, DM, GD and JCMV wrote the manuscript with contributions from all authors.

## Notes

https://github.com/dmoser1/phytocytokine.git

### Competing Interest Statement

The authors have declared no competing interest.

