## Supplementary material for "Maize phytocytokines and microbial-patterns trigger antagonistic features in co-incidence with wounding and fungal pathogens"

**Supplementary Table 1: Organisms represented in BLAST database.** A BLAST database containing 78 predicted plant proteomes was generated. This selection of plant proteomes was randomly chosen from the plants.ensemble.org database and included a variety of monocots, dicots and other phyla.

| **Organisms represented in our customized BLAST database** | | | |
|---|---|---|---|
| *Actinidia chinensis* | *Cynara cardunculus* | *Oryza longistaminata* | *Solanum tuberosum* |
| *Amborella trichopoda* | *Daucus carota* | *Oryza meridionalis* | *Sorghum bicolor* |
| *Ananas comosus* | *Eragrostis curvula* | *Oryza nivara* | *Theobroma cacao criollo* |
| *Arabidopsis halleri* | *Glycine max* | *Oryza punctata* | *Theobroma cacao matina* |
| *Arabidopsis lyrata* | *Gossypium raimondii* | *Oryza rufipogon* | *Trifolium pratense* |
| *Arabidopsis thaliana* | *Helianthus annuus* | *Oryza sativa* | *Triticum aestivum* |
| *Arabis alpina* | *Hordeum vulgare* | *Panicum hallii fil2* | *Triticum aestivum cadenza* |
| *Beta vulgaris* | *Hordeum vulgare goldenpromise* | *Panicum hallii hal2* | *Triticum aestivum claire* |
| *Brachypodium distachyon* | *Ipomoea triloba* | *Phaseolus vulgaris* | *Triticum aestivum paragon* |
| *Brassica napus* | *Lupinus angustifolius* | *Physcomitrella patens* | *Triticum aestivum robigus* |
| *Brassica oleracea* | *Malus domestica golden* | *Pistacia vera* | *Triticum aestivum weebil* |
| *Brassica rapa* | *Medicago truncatula* | *Populus trichocarpa* | *Triticum dicoccoides* |
| *Camelina sativa* | *Nicotiana attenuata* | *Prunus avium* | *Triticum turgidum* |
| *Cannabis sativa female* | *Nymphaea colorata* | *Prunus dulcis* | *Triticum urartu* |
| *Capsicum annuum* | *Olea europaea sylvestris* | *Prunus persica* | *Vigna angularis* |
| *Citrullus lanatus* | *Oryza barthii* | *Rosa chinensis* | *Vigna radiata* |
| *Citrus clementina* | *Oryza brachyantha* | *Selaginella moellendorffii* | *Vitis vinifera* |
| *Coffea canephora* | *Oryza glaberrima* | *Setaria italica* | *Zea mays* |
| *Cucumis melo* | *Oryza glumipatula* | *Setaria viridis* | |
| *Cucumis sativus* | *Oryza indica* | *Solanum lycopersicum* | |

**Supplementary Table 2: Used oligonucleotides**

| ID | Sequence | Use |
|---|---|---|
| GAPDH_F* | CTTCGGCATTGTTGAGGGTTTG | used for qPCR of *gapdh* |
| GAPDH_R* | TCCTTGGCTGAGGGTCCGTC | used for qPCR of *gapdh* |
| ProZip1_F | TGCAGGGAGGAGGGAGATC | used for qPCR of *prozip1* |
| ProZip1_R | CGTCCCTGTTCGGTTCCT | used for qPCR of *prozip1* |
| WYRKY65_F | AGGGCAGTGATTGTCCAAGG | used for qPCR of *wyrky65* |
| WYRKY65_F | TGCTGAAGAGTTGCCGTCTT | used for qPCR of *wyrky65* |
| IGL1_F | GCTGTTGGTTTCGGTGTATCG | used for qPCR of *igl1* |
| IGL1_R | GCCTCACACAACTGCTTCAC | used for qPCR of *igl1* |
| ANK23_F | GGACTTGCAGTTGTTGTCCAG | used for qPCR of *ank23* |
| ANK23_R | AACCCTTCGCTCAGACTTGG | used for qPCR of *ank23* |
| MYC7E_F | AGCACCAGCACCAGAATCAG | used for qPCR of *myc7e* |
| MYC7E_R | CCAAGGAAGGAGTGGACGC | used for qPCR of *myc7e* |
| POX12_F | CTGAACAAGTTCTTCGCGG | used for qPCR of *pox12* |
| POX12_R | AGGTCCACGTAGTACTTGTTG | used for qPCR of *pox12* |
| CC9_F | TATGGGTCCTTGACGTTCTC | used for qPCR of *cc9* |
| CC9_R | GGATCATCCGTAGCCATCTG | used for qPCR of *cc9* |
| PAL1_F | GATCGCCGACATCCAAAGC | used for qPCR of *pal1* |
| PAL1_R | CCTCAAAGAGCACCGTGGAG | used for qPCR of *pal1* |
| PR3_F | GAACAACTACAGCAGCCAGGTG | used for qPCR of *pr3* |
| PR3_R | GAGACAATAGCTGACATGCGTC | used for qPCR of *pr3* |
| PR4_F | GCGTTCAAGCCCATCGACA | used for qPCR of *pr4* |
| PR4_R | CGTGTGGGATCACATCCATATAAC | used for qPCR of *pr4* |
| PR5_F | TATCGGCCGGAATAGGCTCTG | used for qPCR of *pr5* |
| PR5_R | CGCGTACATACAAATGCGTGC | used for qPCR of *pr5* |
| PRm6b_F | CATCTTCGCCATGTTCAACG | used for qPCR of *pr6mb* |
| PRm6b_R | ATTTGTCCGGGTTGAAGAGG | used for qPCR of *pr6mb* |
| PR10.1_F | CAAGCTCATCGCAGACCAC | used for qPCR of *pr10.1* |
| PR10.1_R | CGATCTCAACAGTCCAGCTGTT | used for qPCR of *pr10.1* |

*F: forward (5’-3’), R: reverse (3’-5’)

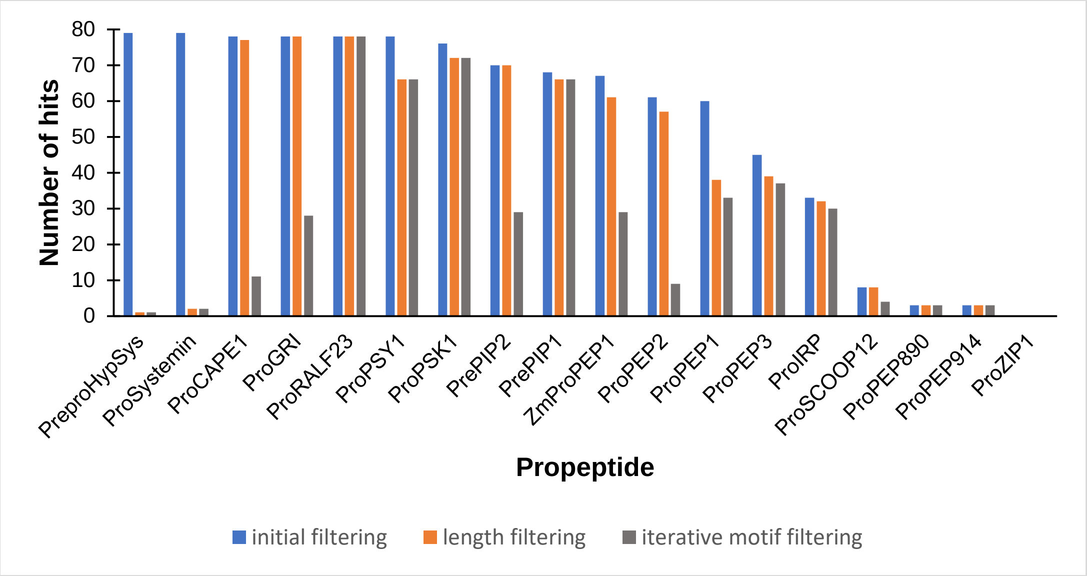


**Supplementary Figure 1. Number of psiBLAST hits after each filtering step.** Unique identifiers were assigned to each psiBLAST hit linking it to the respective propeptide query. Redundant and low-quality hits were removed and only the best hit per organism was selected (initial filtering). Afterwards, hits with low sequence length similiarity to the query (1.5-time longer or shorter) were removed (length filtering). Finally, all hits for a specific query were aligned and the peptide region was identified. The similarity of each peptide homolog to the original peptide was scored by the motif score and hits with motif score below 10% were removed. This step was repeated three times (iterative motif filtering).

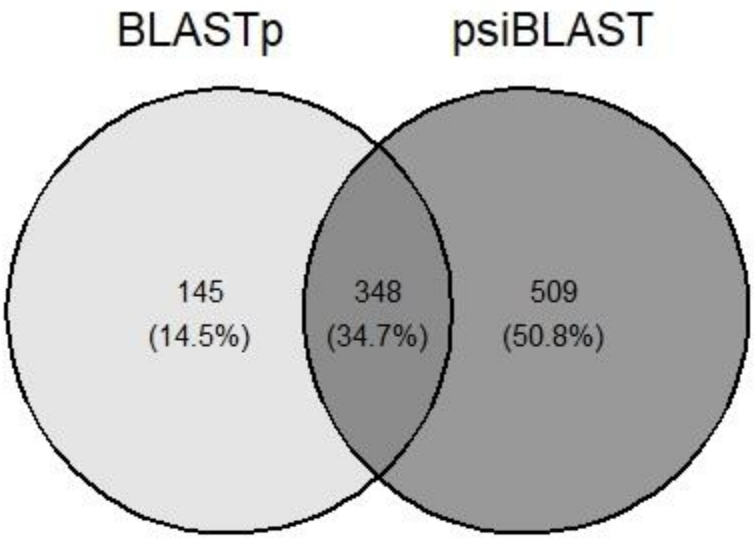


**Supplementary Figure 2. Number of shared and unique hits after BLASTp and psiBLAST.** BLASTp and psiBLAST (Altschul et al., 1997; Camacho et al., 2009) were performed with the phytocytokine protein sequences (Table 1) against a customized BLAST database containing 78 predicted proteomes of selected plant species (Suppl. Table 1). BLASTp and psiBLAST hits were filtered initially by removing redundant and low-quality (protein identities < 10% and query coverage < 25%) hits. Additionally, only the best hit per organism and propeptide based on the bit score was kept. The number of hits found uniquely or shared in the BLASTp or psiBLAST are represented in a venn diagram.

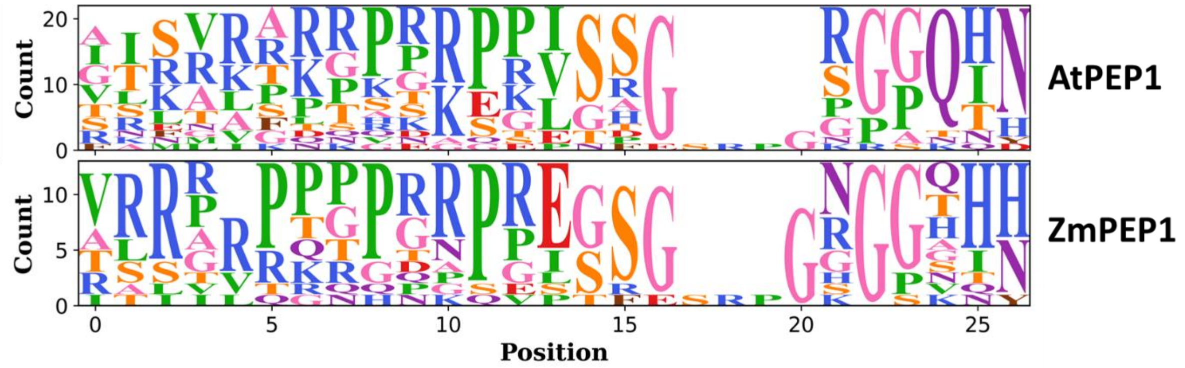


**Supplementary Figure 3. Comparison of across-species peptide motifs derived from AtPEP1 and ZmPEP1 as querys in psiBLAST.** The motifs show a PR-rich N-terminus and a GQHN-rich C-terminus, although with a clear variability. The size of each letter represents the count of a specific amino acid at a given position with position 0 being the first amino acid in the query peptide. Blank spaces indicate gaps in alignments. The functional groups of the amino acids are indicated by color.

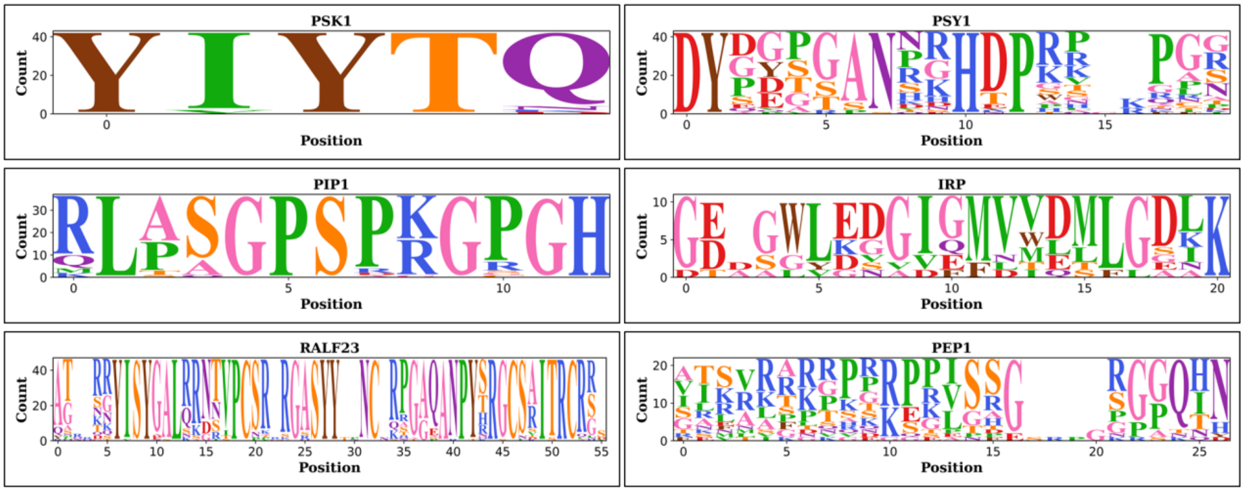


**Supplementary Figure 4. Across-species motifs of the six phytocytokine orthologues found in *Z. mays*.** Sequence similarity of the phytocytokines PSK1, PSY1, PIP1, IRP, RALF23 and PEP1 found in *Z. mays* represented in a motif diagram. The size of each letter represents the count of a specific amino acid at a given position with position 0 being the first amino acid in the query peptide. Blank spaces indicate gaps in alignments. The functional groups of the amino acids are indicated by color.

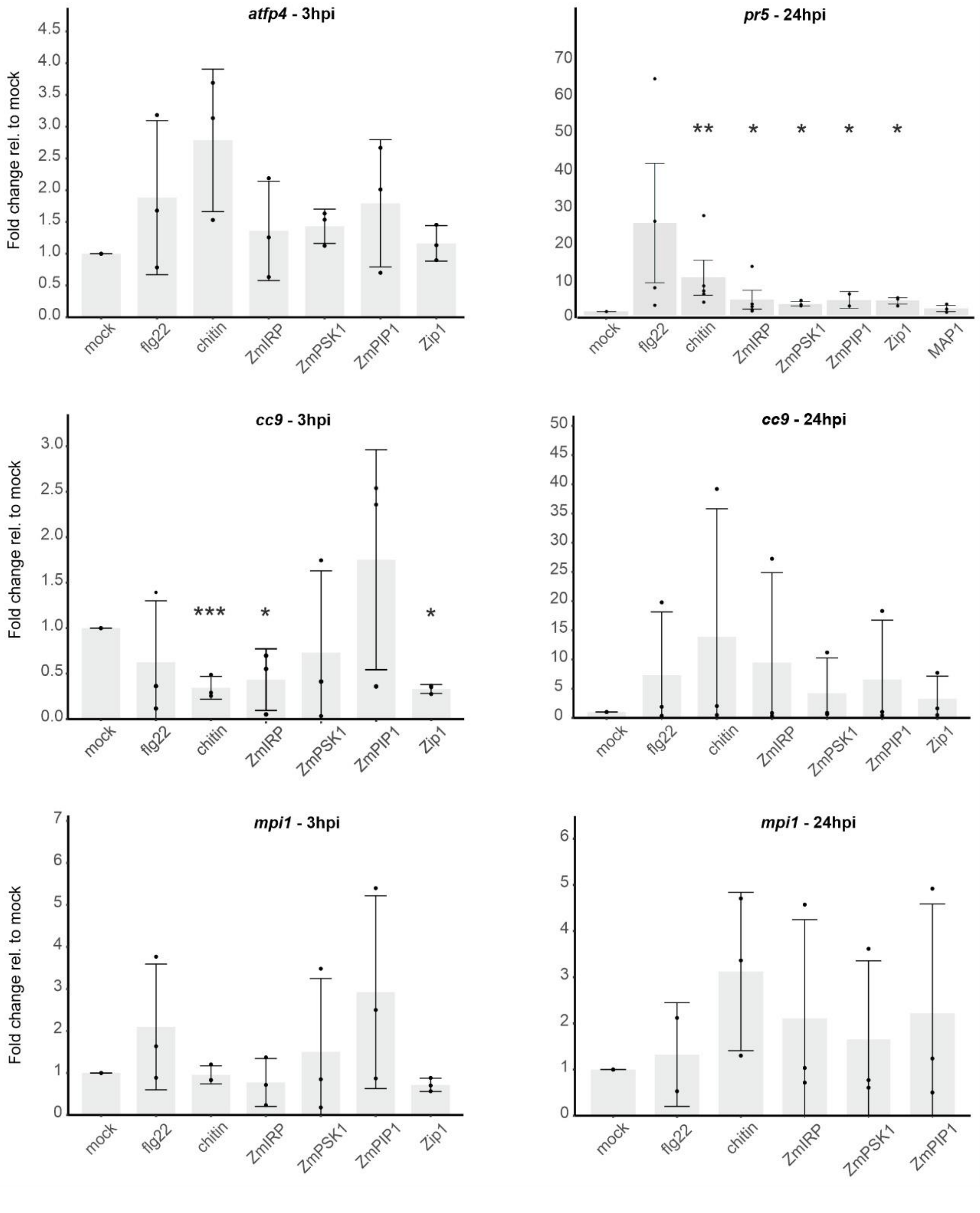


**Supplementary Figure 5. Further marker genes analyzed upon MAMP and phytocytokine treatments.** Shown is the expression of *atfp4* (Hemetsberger et al., 2012), *pr5* (van der Linde et al., 2012), *cc9* (van der Linde et al., 2012) and *mpi1* (Borrego & Kolomiets, 2016) SA- and JA-related marker genes after three and / or 24 hours post infiltration (hpi). Lines show standard error of the mean **(**SEM); p values were calculated via unpaired t-test. *p< 0.05; **p< 0.01; ***p< =0.0001.
